# Hippocampal efferents to retrosplenial cortex and lateral septum are required for memory acquisition

**DOI:** 10.1101/2020.03.23.003996

**Authors:** Ashley N Opalka, Dong V Wang

## Abstract

Learning and memory involves a large neural network of many brain regions, including the notable hippocampus along with the retrosplenial cortex (RSC) and lateral septum (LS). Previous studies have established that the dorsal hippocampus (dHPC) plays a critical role during the acquisition and expression of episodic memories. However, the role of downstream circuitry from the dHPC, including the dHPC-to-RSC and dHPC-to-LS pathways, has come under scrutiny only recently. Here, we employed an optogenetic approach with contextual fear conditioning in mice to determine whether the above two pathways are involved in acquisition and expression of contextual fear memory. We found that a selective inhibition of the dHPC neuronal terminals in either the RSC or LS during acquisition impaired subsequent memory performance, suggesting that both the dHPC-to-RSC and dHPC-to-LS pathways play a critical role in memory acquisition. We also selectively inhibited the two dHPC efferent pathways during memory expression and found a differential effect on memory performance. These results indicate the intricacies of memory processing and that hippocampal efferents to cortical and subcortical regions may be differentially involved in aspects of physiological and cognitive memory processes.

## Introduction

Processing of memory entails multiple stages: acquisition, the initial encoding of memory; consolidation, the transformation of newly acquired information into long-lasting memory; retrieval, the recall of stored memory; and expression, the behavioral readout of recalled memory. Extensive research has shown that the dorsal hippocampus (dHPC) plays an important role in each of these memory processing stages (Scoville and Milner, 1957, Maren and Fanselow, 1997, McEchron et al., 1998, Lee and Kesner, 2004, Misane et al., 2005, Carr et al., 2011, Goshen et al., 2011, Pierson et al., 2015, Ocampo et al., 2017). However, how the dHPC communicates with downstream cortical and subcortical regions during these memory stages remains unclear. Recent studies have shown that a number of cortical regions display increased activity during memory acquisition and expression stages (Takata et al., 2015, Miller et al., 2017, DeNardo et al., 2019), and that hippocampal-cortical communications are required for memory acquisition and consolidation (Frankland et al., 2004, Kitamura et al., 2017, DeNardo et al., 2019, Yamawaki et al., 2019a). Thus, recruitment of cortical brain regions during the initial acquisition stage may be necessary for subsequent consolidation and retrieval/expression of memories (Lesburguères et al., 2011, Kitamura et al., 2017, DeNardo et al., 2019, de Sousa et al., 2019). Additionally, recruitment of hippocampal projections to subcortical regions appears to be necessary during these memory processes as well (Olton et al., 1978, Hunsaker et al., 2009, Roy et al., 2017, Besnard et al., 2019, Besnard et al., 2020).

The dHPC projects directly to a few cortical and subcortical regions, notably the granular retrosplenial cortex (RSC) and the lateral septum (LS) (Van Groen and Wyss, 1990, Jinno et al., 2007, Miyashita and Rockland, 2007, Kwapis et al., 2015, Takata et al., 2015, Todd and Bucci, 2015). However, the function of these two dHPC projections during memory processes has come under scrutiny only recently (Yamawaki et al., 2019a, Yamawaki et al., 2019b, Opalka et al., 2020, Besnard et al., 2020). Previous studies revealed that the RSC contributes to contextual memory (Keene and Bucci, 2008, Corcoran et al., 2011, Cowansage et al., 2014, Kwapis et al., 2015, Robinson et al., 2018, de Sousa et al., 2019), spatial memory (Cooper et al., 2001, Vann and Aggleton, 2002, Czajkowski et al., 2014, Mao et al., 2017, Milczarek et al., 2018), inhibitory avoidance memory (Katche et al., 2013, Katche and Medina, 2017), and multi-sensory association (Robinson et al., 2011, Robinson et al., 2014). Moreover, the RSC is one of few cortical regions that receive direct projections from the dHPC, positioning the RSC as a potential bridge in connecting the hippocampus with other cortical regions, such as the anterior cingulate cortex, medial prefrontal cortex and secondary sensory cortices, to support long-term memory (Maviel et al., 2004, Frankland et al., 2004, Todd and Bucci, 2015, Wang and Ikemoto, 2016, Kitamura et al., 2017, DeNardo et al., 2019). Therefore, a better understanding of this dHPC-to-RSC connection will shed light on memory acquisition and retrieval/expression processes.

Further, previous literature reports the LS as essential for processing multiple forms of memory, such as contextual memory (Calandreau et al., 2007, Calandreau et al., 2010, Besnard et al., 2019, Besnard et al., 2020), social encounter and memory (Dantzer et al., 1988, Everts and Koolhaas, 1997, Leroy et al., 2018), and addiction memory (McGlinchey and Aston-Jones, 2018). Due to the hippocampal-septal involvement in theta oscillation (Colgin, 2016), the dHPC-to-LS pathway has been recently investigated for its role in spatial navigation and spatial memory (Tingley and Buzsáki, 2018). Specifically, the firing of LS neurons phase-locked to dHPC theta oscillation (Mondragón-Rodríguez et al., 2019) positions this dHPC-to-LS pathway as a candidate for memory processing. To test whether the dHPC-to-RSC and dHPC-to-LS neural pathways are involved in the acquisition and retrieval/expression of memory, we employed an optogenetic approach and contextual fear conditioning, a procedure widely used to access hippocampus-dependent memory. Our results provide direct evidence that both the cortical (RSC) and subcortical (LS) projections from the dorsal hippocampus are required during acquisition of the fear memory. These hippocampal efferents to downstream cortical and subcortical regions may process distinct features in representing a memory in its entirety.

## Results

We bilaterally injected AAV viruses that encode fluorescent eYFP under the promoter CaMKII into the medial portion of the dHPC region (including medial CA1 and dorsal subiculum; Suppl. figures 1 and 2). Consistent with previous work (Wyss and Van Groen, 1992, Oh et al., 2014), our results showed that this dHPC region projected primarily to the granular RSC layer 3, midline LS, entorhinal cortex, and mammillary area. In the present study, we optogenetically inhibited two dHPC efferent pathways: the dHPC-to-RSC and dHPC-to-LS pathways (Suppl. figure 1). Repeated measures two-way ANOVAs were conducted on freezing for each of the following experiments followed by Bonferroni post-hoc analyses. The between-subjects variables included two levels of treatment: halorhodopsin (Halo) or eYFP (Ctrl) viral injections. The within-subjects variables included two levels of time: recent (day 1) or remote (day 31) memory tests.

### Optogenetic inhibition of dHPC-to-RSC pathway during memory acquisition

Mice received bilateral dHPC injections, targeting medial CA1 and rostral-dorsal subiculum (Suppl. figures 1 and 2), of either AAV-Halo (Halo) or AAV-eYFP (Ctrl). Meanwhile, an optical fiber was implanted in the midline of the RSC, capable of inhibiting dHPC projections from both hemispheres because the dHPC projects predominantly to RSC layer 3, located close to the midline (Figure 1A, left; Suppl. figure 1). Four-five weeks after surgery, mice received photoinhibition of the dHPC-to-RSC neuronal terminals during acquisition of contextual fear memory on day 0. Mice were tested subsequently for recent memory in 1 day and remote memory in 31 days, receiving no photoinhibition during the tests (Figure 1A, right). There was a significant main effect of viral group (F_1,46_ = 9.384, *P* = 0.004; repeated measures two-way ANOVA; Figure 1B), indicating a substantial and enduring memory deficit of the Halo group compared to the Ctrl group. In addition, there was a main effect of time (F_1,46_ = 6.317, *P* = 0.016), indicating a difference in freezing between recent and remote memory tests, but there was no interaction effect (F_1,46_ = 0.266, *P* = 0.608). Further minute-by-minute data of freezing are shown in Figure 1C. These results suggest that the dHPC-to-RSC neural pathway plays a critical role in acquisition of contextual fear memory. Note that Figure 1B & 1C combined two datasets (n = 24 per dataset, n = 12 per group; see Methods for details and Suppl. figure 3 for data/statistics).

**Figure 1.**
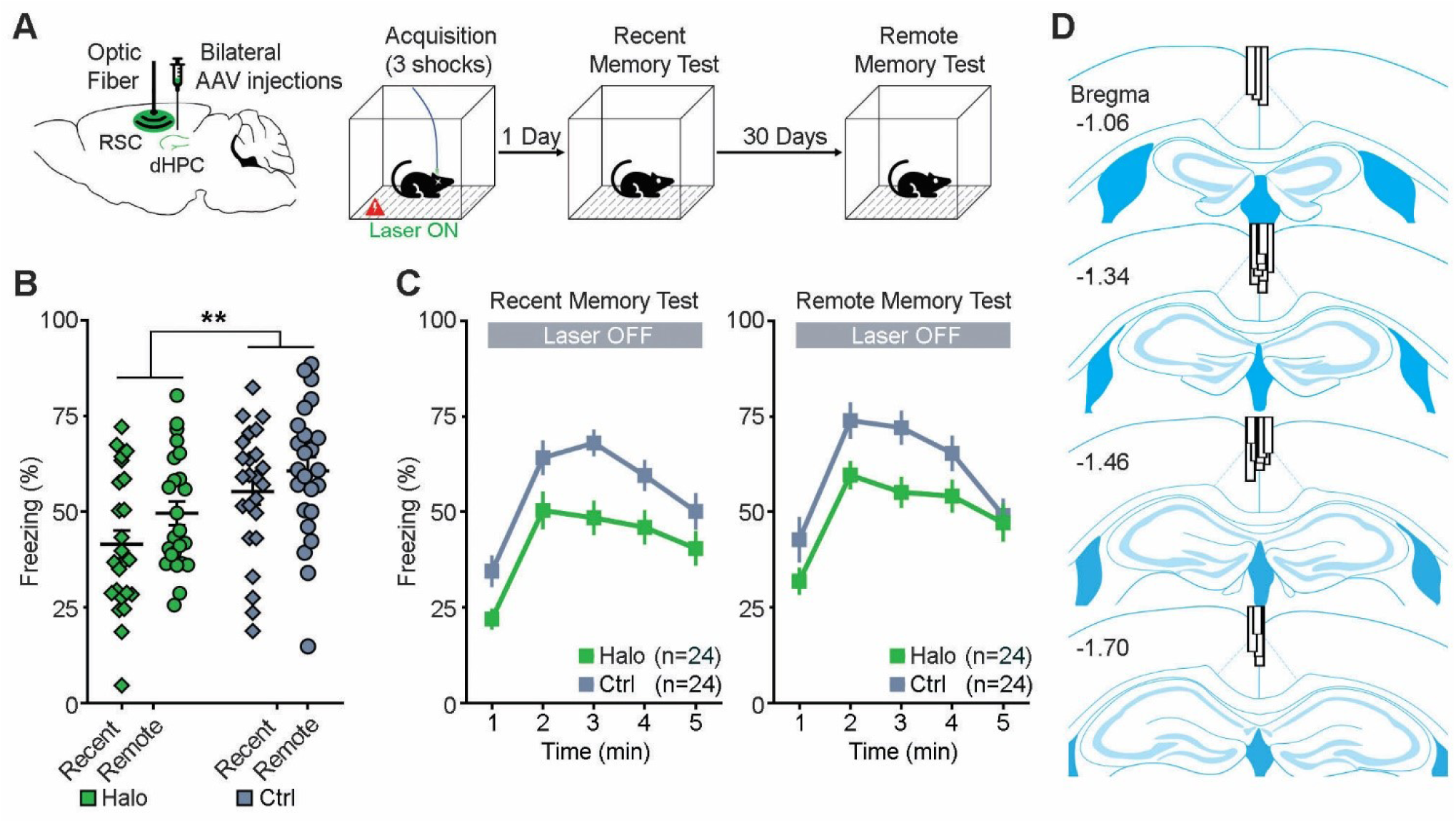
Optogenetic inhibition of dHPC-to-RSC pathway during memory acquisition. (A) Schematic drawing (left) of bilateral injections of AAV viruses that encode either halorhodopsin (Halo) or YFP (Ctrl) into the dHPC and an optical fiber implant into the RSC. Contextual fear conditioning procedure (right). Mice received optogenetic inhibition of the dHPC-to-RSC neuronal terminals during acquisition of memory on day 0. Then mice were tested for recent memory on day 1 and remote memory on day 31, receiving no laser stimulation. (B) Percentage of freezing (mean of 5 minutes) during the recent memory test (day 1) and remote memory test (day 31). ** *P* = 0.004, repeated measures ANOVA between groups. (C) Minute-by-minute percentage of freezing from B of recent (left) and remote (right) memory tests. Both left & right, *P* < 0.05; Bonferroni post hoc of the means between halo/Ctrl. All error bars indicate standard error of the mean (SEM). (D) Optical fiber placements of the Halo group were confirmed mostly dorsal to the RSC, arranged anterior (top) to posterior (bottom). Adapted from Franklin and Paxinos 2008.

**Figure 2.**
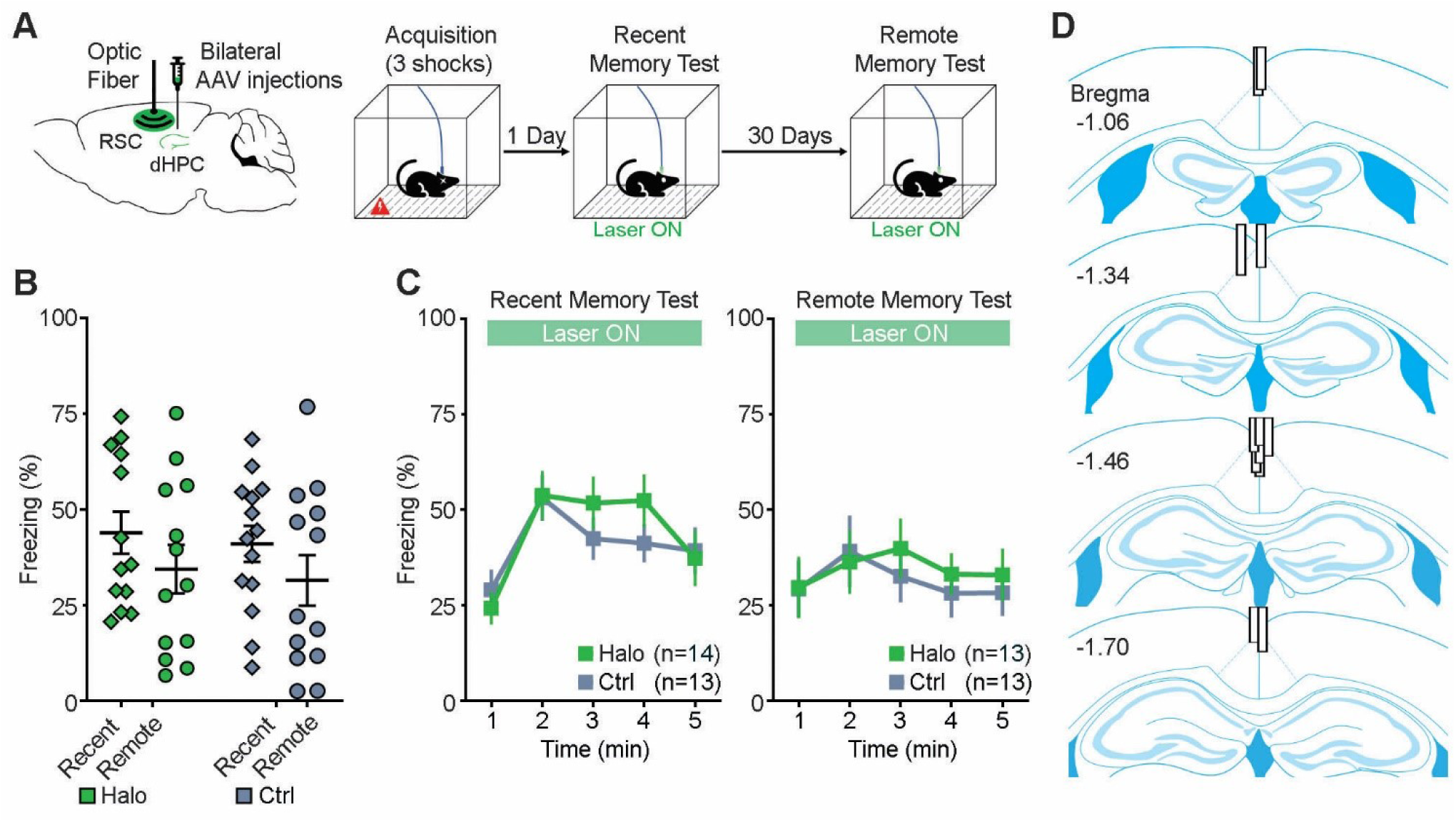
Optogenetic inhibition of dHPC-to-RSC pathway during memory retrieval. (A) Schematic drawing (left) of bilateral injections of AAV viruses that encode either halorhodopsin (Halo) or YFP (Ctrl) into the dHPC and an optical fiber implant into the RSC. Contextual fear conditioning procedure (right). Mice were trained during acquisition of memory on day 0, receiving no laser stimulation. Then, mice received optogenetic inhibition of the dHPC-to-RSC neuronal terminals during both recent (day 1) and remote (day 31) memory tests. (B) Percentage of freezing (mean of 5 minutes) during the recent memory test (day 1) and remote memory test (day 31). *P* = 0.694; repeated measures ANOVA between groups. (C) Minute-by-minute percentage of freezing from B of recent (left) and remote (right) memory tests. All error bars indicate SEM. (D) Optical fiber placements of the Halo group were confirmed mostly dorsal to the RSC, arranged anterior (top) to posterior (bottom).

On the day of training (memory acquisition; day 0), we also analyzed freezing and, additionally, motion index to determine if the laser stimulation impacted locomotion. We found that both freezing during the total training session (*t*_(46)_ = -0.091, *P* = 0.928; unpaired *t* test; Suppl. figure 4A, top left) and motion index during the first two minutes before footshocks (*t*_(46)_ = -0.207, *P* = 0.837; Suppl. figure 4A, top right) did not differ between the Halo and Ctrl groups. This suggests that the laser stimulation did not affect freezing expression or general locomotion. After completion of the above experiments, histological verification of optical fiber placements and viral injections revealed that all optical fiber implants were chronically placed around the midline and dorsal to the RSC (Figure 1D; Suppl. figure 1B), and all dHPC expressed fluorescence. Therefore, all animals were included for analyses.

### Optogenetic inhibition of dHPC-to-RSC pathway during memory retrieval

Similarly, mice received bilateral dHPC injections of either AAV-Halo or AAV-eYFP and a RSC optical fiber implantation (Figure 2A, left). Four-five weeks after surgery, mice were trained for acquisition of contextual fear memory on day 0 and were tested subsequently for recent (day 1) and remote (day 31) memories. Mice received photoinhibition of the dHPC-to-RSC neuronal terminals during both testing days (Figure 2A, right). There was no main effect of group (F_1,24_ = 0.159, *P* = 0.694; repeated measures two-way ANOVA; Figure 2B) or interaction effect (F_(1,46)_ = 0.001, *P* = 0.973), but a main effect of time (F_1,24_ = 7.053, *P* = 0.014), indicating a difference in freezing between recent and remote memory tests. Further minute-by-minute data of freezing are shown in Figure 2C. In addition, there was no significant difference in mean freezing between Halo and Ctrl viral groups during acquisition (day 0; *t*_(46)_ = 0.648, P = 0.523; unpaired *t* test; Suppl. figure 4B, top). These results suggest that the dHPC-to-RSC neural pathway is dispensable during the retrieval/expression of contextual fear memory, though we cannot exclude the possibility of an insufficient inhibition in our experiment. After experiments concluded, histological verification revealed that all optical fiber implants were chronically placed near the midline and dorsal to the RSC (Figure 2D; Suppl. figure 1B) and that all dHPC expressed fluorescence. All animals were included for analyses expect one (Ctrl) that lost its optical fiber implant after the recent memory test and was sacrificed before the remote memory test.

### Optogenetic inhibition of dHPC-to-LS pathway during memory acquisition

Mice received bilateral dHPC injections of AAV-Halo or AAV-eYFP and an optical fiber implant into the midline immediately above the LS (Figure 3A, left). This implant is capable of bilateral inhibition because the dHPC (our injection site) projects mainly to the midline rostral LS (Suppl. figure 1). Four-five weeks after surgery, mice received photoinhibition of the dHPC-to-LS neuronal terminals during acquisition of contextual fear memory on day 0. Mice were tested subsequently for recent (day 1) and remote (day 31) memories, receiving no photoinhibition during the tests (Figure 3A, right). There was a significant main effect of viral group (F_1,24_ = 4.623, *P* = 0.042; repeated measures two-way ANOVA; Figure 3B), indicating a substantial and enduring memory deficit in the Halo group compared to the Ctrl group. There was a trending difference on main effect of time (F_1,24_ = 4.066, *P* = 0.055) but no interaction effect (F_1,24_ = 0.011, *P* = 0.917). Further minute-by-minute data of freezing are shown in Figure 3C. Together, these results suggest that the dHPC-to-LS neural pathway plays a critical role in the acquisition of contextual fear memory.

**Figure 3.**
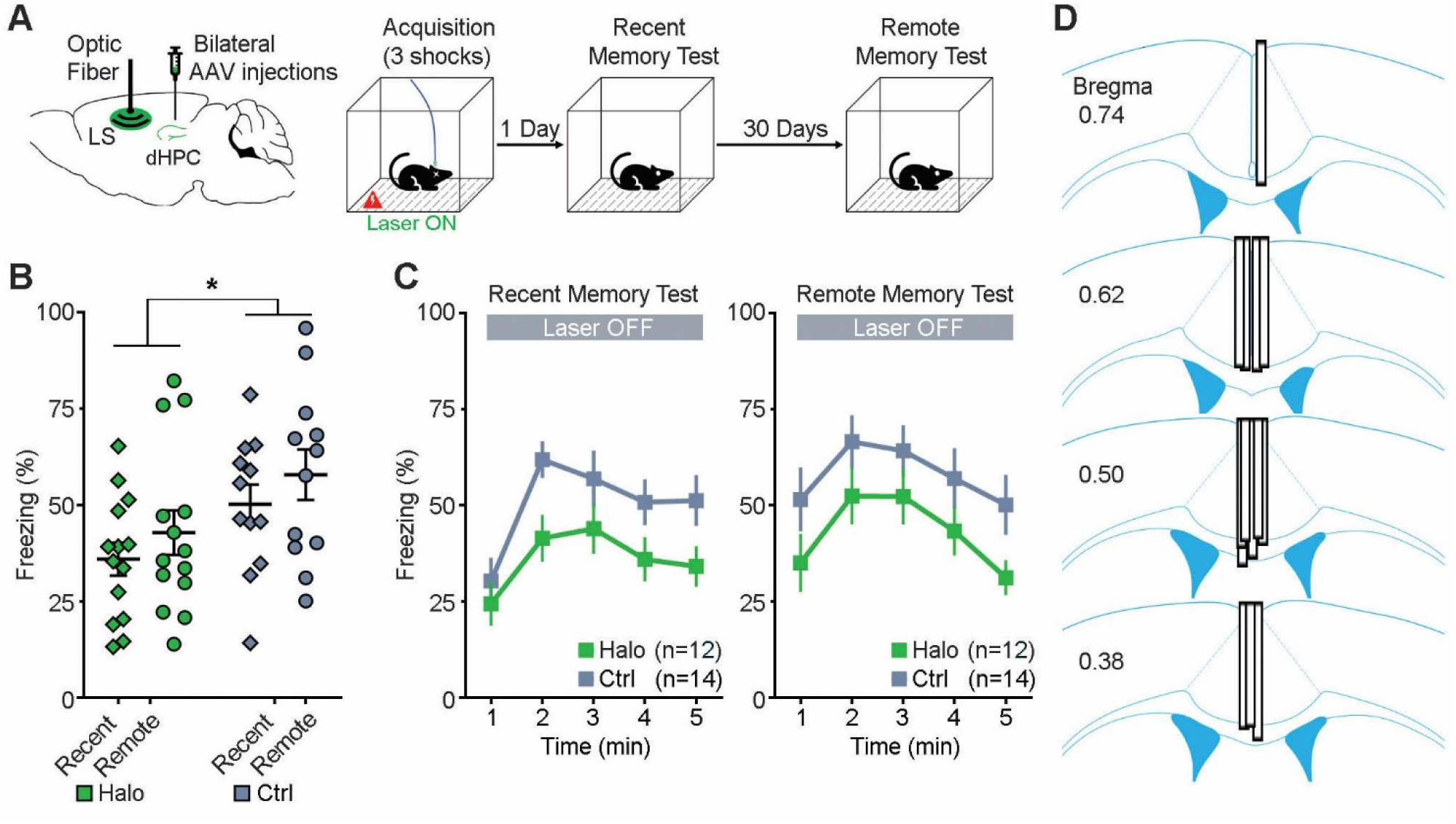
Optogenetic inhibition of dHPC-to-LS pathway during memory acquisition. (A) Schematic drawing (left) of bilateral injections of AAV viruses that encode either halorhodopsin (Halo) or YFP (Ctrl) into the dHPC and an optical fiber implant into the LS. Contextual fear conditioning procedure (right). Mice received optogenetic inhibition of the dHPC-to-LS neuronal terminals during acquisition of memory on day 0. Then were tested for recent memory on day 1 and remote memory on day 31, receiving no laser stimulation. (B) Percentage of freezing (mean of 5 minutes) during the recent memory test (day 1) and remote memory test (day 31). * *P* = 0.042; repeated measures ANOVA between groups. (C) Minute-by-minute percentage of freezing from B of recent (left) and remote (right) memory tests. Both left & right, *P* > 0.05; Bonferroni post hoc of the means between halo/Ctrl. All error bars indicate SEM. (D) Optical fiber placements of the Halo group were confirmed mostly dorsal to the LS, arranged anterior (top) to posterior (bottom).

We also analyzed the freezing during training (memory acquisition) on day 0. Optogenetic inhibition of the dHPC-to-LS pathway during the total training session did not impact freezing between Ctrl and Halo groups (*t* _(24)_ = 1.306, *P* = 0.218; unpaired *t* test; Suppl. figure 4A, bottom left), suggesting that the dHPC-to-LS pathway does not impair freezing expression. Furthermore, we analyzed motion index during training (memory acquisition, day 0) to determine if laser stimulation altered locomotion, which trended toward significance during the first two minutes before footshocks (*t* _(24)_ = -2.040, *P* = 0.053; unpaired *t* test; Suppl. figure 4A, bottom right). To rule out any locomotor effect, we further conducted a locomotor test with a subset of mice and again found a trending difference between Halo (n = 8) and Ctrl (n = 7) groups upon laser stimulation (*t*_(13)_ = -1.945, *P* = 0.074; unpaired *t* test; see Suppl. Methods). This suggests that optogenetic inhibition of the dHPC-to-LS pathway has no or limited effect on locomotor activity. After completion of the above experiments, histological verification revealed that all optical fibers were chronically placed around the midline and dorsal to the rostral LS (Figure 3D; Suppl. figure 1C), and all dHPC expressed fluorescence. Therefore, all animals were included for analyses.

### Optogenetic inhibition of dHPC-to-LS pathway during memory retrieval

Similarly, mice received bilateral dHPC injections of either AAV-Halo or AAV-eYFP and a LS optical fiber implantation (Figure 4A, left). Four-five weeks after surgery, mice were trained for acquisition of contextual fear memory on day 0 and were tested subsequently for recent memory on day 1 and remote memory on day 31. Mice received photoinhibition of the dHPC-to-LS neuronal terminals during both testing days (Figure 4A, right). There was a significant main effect of viral group (F_1,25_ = 9.818, *P* = 0.004; repeated measures two-way ANOVA; Figure 4B), indicating impaired memory of the Halo group compared to the Ctrl group. There was no main effect of time (F_1,25_ = 2.363, *P* = 0.137) or interaction effect (F_1,25_ = 0.933, *P* = 0.343). Further minute-by-minute data of freezing are shown in Figure 4C. These results suggest that the dHPC-to-LS neural pathway plays a critical role in the expression of contextual fear memory. No significant difference of freezing existed between groups during acquisition of memory (day 0; *t* _(25)_ = 0.943, *P* = 0.335; unpaired *t* test; Suppl. figure 4B, bottom). After experiments concluded, histological verification revealed that all optical fibers were chronically placed on the midline and dorsal to the LS (Figure 4D; Suppl. figure 1C) and that all dHPC expressed fluorescence. Therefore, all animals were included for analyses.

**Figure 4.**
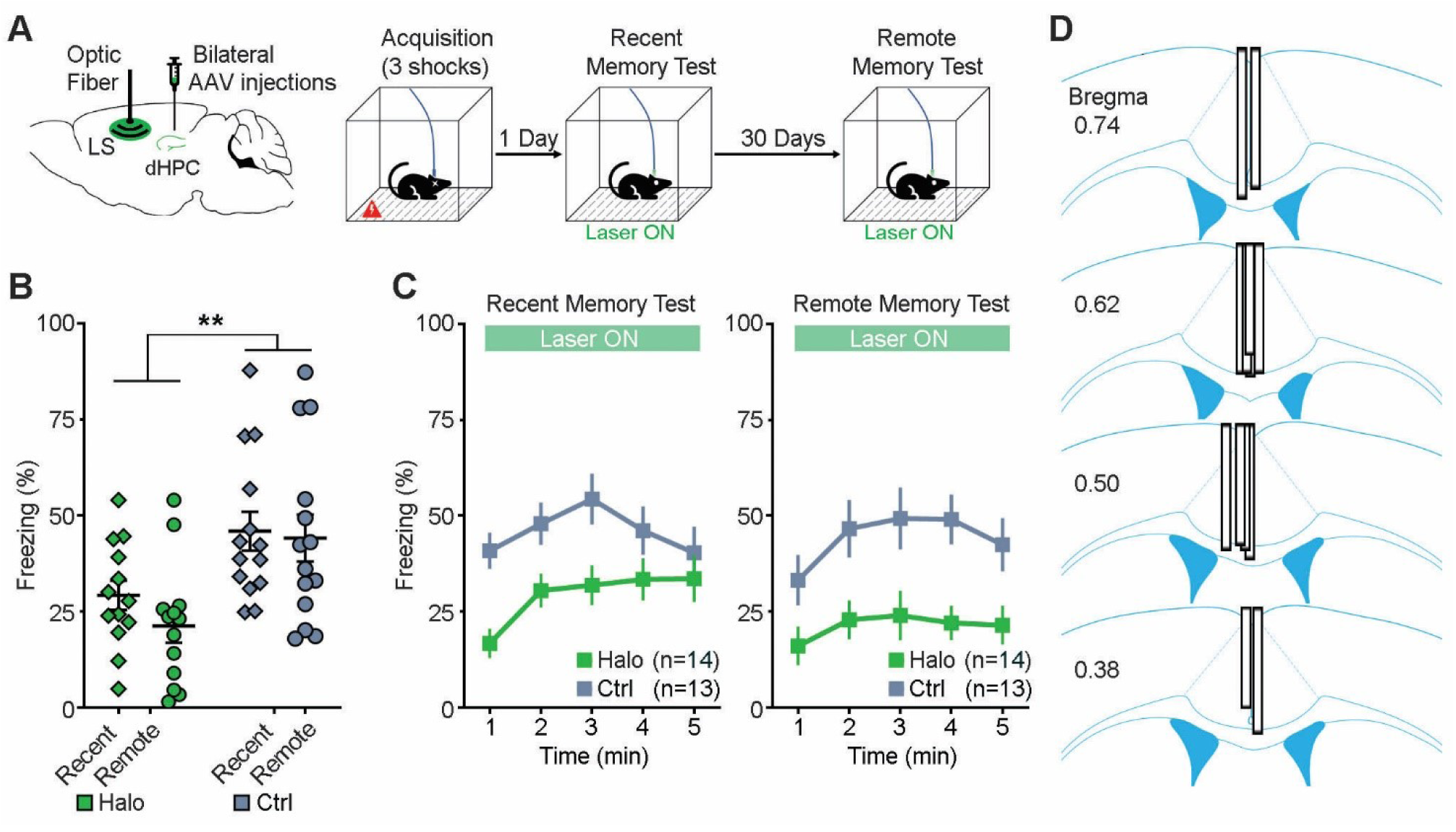
Optogenetic inhibition of dHPC-to-LS pathway during memory retrieval. (A) Schematic drawing (left) of bilateral injections of AAV viruses that encode either halorhodopsin (Halo) or YFP (Ctrl) into the dHPC and an optical fiber implant into the LS. Contextual fear conditioning procedure (right). Mice were trained during acquisition of memory on day 0, receiving no laser stimulation. Then, mice received optogenetic inhibition of the dHPC-to-LS neuronal terminals during both memory recent (day 1) and remote (day 31) memory tests. (B) Percentage of freezing (mean of 5 minutes) during the recent memory test (day 1) and remote memory test (day 31). ** *P* = 0.004; repeated measures ANOVA between groups. (C) Minute-by-minute percentage of freezing from B of recent (left) and remote (right) memory tests. Both left & right, *P* < 0.05; Bonferroni post hoc of the means between halo/Ctrl. All error bars indicate SEM. (D) Optical fiber placements of the Halo group were confirmed mostly dorsal to the LS, arranged anterior (top) to posterior (bottom).

### Optogenetic inhibition of dHPC terminals induces firing changes in the RSC and LS

To confirm that the dHPC projections can influence downstream neural activity, we utilized a combined optogenetic and electrophysiology approach (Figure 5): we unilaterally injected AAV-Halo viruses into the dHPC and implanted an optrode (an optical fiber attached to eight tetrodes) into the ipsilateral granular RSC or LS. Three-four weeks after the surgery, we recorded RSC or LS spike activity in freely behaving mice while intermittently delivering photoinhibitions (5 min in every 15 min; ∼2 h per session). Our results revealed that photoinhibition of the dHPC projections affected firings of both RSC and LS neurons. A subpopulation of the RSC and LS neurons was silenced or suppressed, demonstrating the effectiveness of our photoinhibition approach (Figure 5 B,C,E,F). Interestingly, another subpopulation of the RSC and LS neurons were activated upon the same photoinhibition (Figure 5 B,C,E,F), which could be explained by a disinhibition mechanism via local interneurons (Opalka et al., 2020) or a network feedback disinhibition. Notably, these results also revealed a functional recovery of the RSC and LS neurons, albeit a small rebound activation in a subset of neurons after the cessation of the photoinhibition (Figure 5 B,C,E,F).

**Figure 5.**
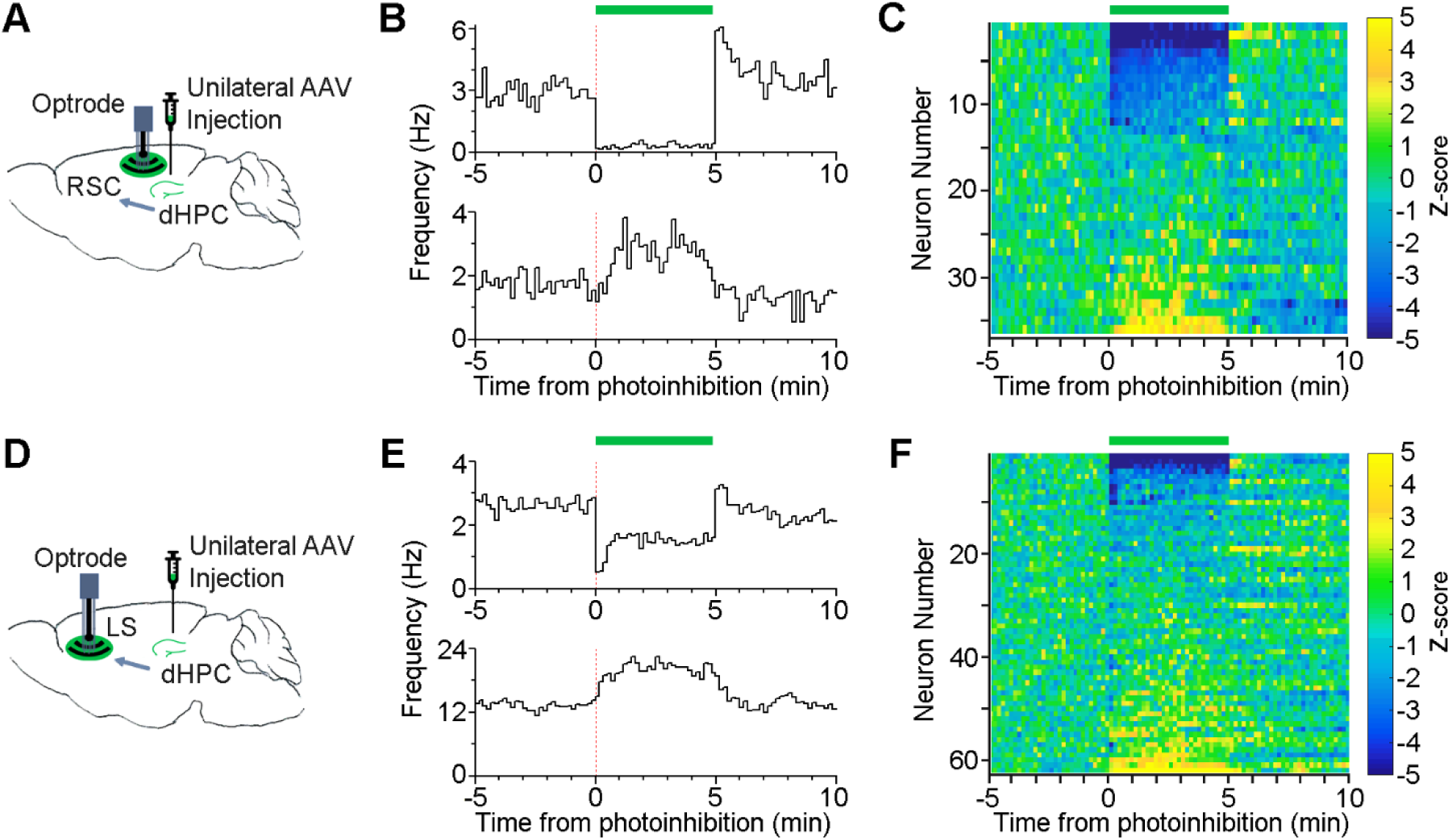
Optogenetic inhibition of dHPC terminals induces RSC and LS firing changes. (A&D) Schematic drawing of unilateral viral injection of AAV-CaMKII-Halo into the dHPC and an optrode implant, an optical fiber attached to eight tetrodes, into either the RSC (A) or LS (D). (B&E) Two example RSC (B) or LS (E) neurons recorded simultaneously that were inhibited (top) or activated (bottom) upon photoinhibition (green bar, 0–5 min; bin, 10 s). (C&F) Heatmap of all RSC (C) and LS (F) neurons upon photoinhibition (green bar, 0–5 min). Color bars, z-scored firing rate.

## Discussion

Utilizing an optogenetic approach and contextual fear conditioning paradigm, we provided evidence that the dHPC-to-RSC neural pathway is critical for memory acquisition, whereas the dHPC-to-LS neural pathway is critical for both memory acquisition and expression. In both instances, the memory impairments were similar between recent and remote memory tests (one month apart), indicating the reliability of optogenetic manipulation on memory acquisition and expression. Notably, this optogenetic approach enabled transient and reversible inhibition of hippocampal efferent pathways compared to lesions or drug infusions that often cause permanent damages or long-lasting neural alterations. Additionally, our use of the CaMKII promotor likely only targeted hippocampal pyramidal neurons but not interneurons (Stark et al., 2012, Wang et al., 2013, Zhu et al., 2014). With our optogenetic approach, it is noted that there’s limited spread of AAV viruses to the dentate gyrus, which does not project to the RSC or LS (Van Groen and Wyss, 1990, Risold and Swanson, 1997, Witter, 2007). We also observed AAV leak in cortical regions along the injection needle track, in some cases spreading to ventral lateral RSC near the corpus callosum (Suppl. figure 2). These leaks were relatively small but could be partially responsible for our observed behavioral changes. On the other hand, we did not see any AAV spread to the LS, which is expected given the long distance between the injection site (medial CA1) and LS. Together, our observed memory deficits should be attributed primarily to the dHPC-to-RSC or dHPC-to-LS excitatory pathway and its functions.

### dHPC-to-RSC pathway is critical for memory acquisition

Previous studies have shown that the RSC plays a critical role during acquisition of multiple forms of memories, including acquisition of contextual fear memory (Corcoran et al., 2011, Cowansage et al., 2014, Kwapis et al., 2015, Todd et al., 2016, de Sousa et al., 2019), inhibitory avoidance memory (Katche and Medina, 2017) and trace fear memory (Kwapis et al., 2015). Moreover, employing chemogenetic methods, two recent studies found that the dHPC-to-RSC excitatory and inhibitory projections play opposing roles in facilitating and inhibiting memory acquisition, respectively (Yamawaki et al., 2019a, Yamawaki et al., 2019b). Consistently, by employing optogenetic methods, we provided evidence that the dHPC-to-RSC projection plays a critical role during memory acquisition. In contrast, the entorhinal-to-RSC pathway seems to be dispensable during memory acquisition memories (Kitamura et al., 2017), indicating a functional differentiation of RSC connections. Together, our results add to the growing evidence that hippocampal-cortical communication during memory acquisition is critical for the formation of both recent and remote memories (Kitamura et al., 2017, DeNardo et al., 2019).

Correspondingly, the RSC has been shown to play a critical role during retrieval/expression of multiple forms of memories, including contextual fear memory (Corcoran et al., 2011, Cowansage et al., 2014, Kwapis et al., 2015, Todd et al., 2016), inhibitory avoidance memory (Katche et al., 2013), cued fear memory (Todd et al., 2016) and autobiographical memory (Svoboda et al., 2006). However, it remains unclear which upstream regions control the RSC and support memory retrieval/expression, given that the RSC receives direct inputs from a number of memory associated regions, such as the dHPC, thalamus and anterior cingulate cortex (Todd et al., 2016). A recent study reported that chemogenetic inhibition of dHPC glutamatergic (VGLUT1) terminals in the RSC impaired the expression of recent but not remote memory (Yamawaki et al., 2019a), whereas our optogenetic inhibition of the dHPC-to-RSC pathway had little effect on the expression of recent or remote memories (albeit a potential insufficient inhibition). One justification for this discrepancy could be due to spatial restriction and temporal sensitivity of the optogenetic approach compared to chemogenetics. This discrepancy may also be due to different volume viral injections: we injected 0.2 µL per hemisphere, limiting viral expression primarily to the medial (distal) CA1 and rostral-dorsal subiculum, whereas their larger injection of 0.5 µL per hemisphere likely infected a broader dHPC area, such as the lateral (proximal) CA1 and caudal subiculum. These alternative lateral CA1 and caudal subiculum efferents may support memory retrieval, whereas the medial CA1 and rostral subiculum efferents may support memory acquisition (Roy et al., 2017, Nakazawa et al., 2016).

### dHPC-to-LS pathway is critical for memory acquisition and expression

We found that transient optogenetic inhibition of dHPC projections to the LS during contextual fear acquisition hindered subsequent memory performances, indicating the importance of the dHPC-to-LS pathway in memory acquisition. This is consistent with a previous study showing that inactivation of the LS impaired contextual fear acquisition (Calandreau et al., 2007). Additionally, we found that optogenetic inhibition of dHPC projections to the LS during contextual fear retrieval/expression disrupted fear memory performance, indicating the importance of this dHPC-to-LS pathway in memory expression as well, corroborating with earlier studies (Olton et al., 1978, Besnard et al., 2020). Together, our results suggest that the dHPC-to-LS pathway is critical for both memory acquisition and expression. In support of this, a previous study showed that transected hippocampal efferents to the subcortex impaired both the acquisition and expression of memory (Hunsaker et al., 2009).

One caveat regards the potential photoinhibition of LS passing fibers (from the dHPC) to additional downstream subcortical regions, such as the diagonal band of Broca and mammillary regions. Future experiments that specifically target these dHPC projections are needed to understand their functions. Additionally, our observed deficit during memory expression may be confounded by reduced freezing expression because the dHPC-to-LS and other LS input pathways have been implicated in processing animal’s locomotor speed (Bender et al., 2015, Wirtshafter and Wilson, 2019), despite that manipulating the CA1 itself does not affect locomotion (Goshen et al., 2011). In our experiments, inhibition of the dHPC-to-LS pathway did not affect freezing during training but slightly increased locomotion (Suppl. figure 4). Therefore, additional experiments, that do not rely on freezing as a measurement of memory, are necessary to validate the functional role of this dHPC-to-LS pathway in memory retrieval/expression in the future.

### Recent and remote memories

Our results showed that the memory impairments were similar between recent and remote memory tests (one month apart). Notably, there was a slight increase of freezing between recent and remote memory tests for mice without optical fiber attachment (Figure 1B; Figure 3B). This is consistent with previous studies that reported strengthened remote memory compared to recent memory (Broadbent et al., 2006, Frankland et al., 2004, Blum et al., 2006). However, there was a decrease of freezing between recent and remote memory tests for mice that had an optical fiber attachment (Figure 2B; Figure 4B). This reattachment of the optical fiber during remote memory tests may have distracted the mice and affected their expression of freezing, given that the mice had not experienced the fiber attachment for one month. Nonetheless, the fiber attachment seemed to have little effect on the recent memory test because animals were habituated to the fiber attachment 24 h prior to the test.

Together, our study along with others reveal the intricate functional selectivity of memory pathways and provide evidence that multiple memory systems, involving select cortical and subcortical regions during varying stages of memory, may activate in parallel or, conversely, compete under certain circumstances. The dorsal hippocampal efferents to both cortical (RSC) and subcortical (LS) regions play an important role in memory acquisition. This similar role of the dHPC-to-RSC and dHPC-to-LS projections in fear memory acquisition suggests that the two parallel pathways may contribute to the same or represent different aspects of the fear memory, such as freezing, stress hormones, blood pressure, etc. (Roy et al., 2017). Disruption of these pathways during acquisition may also prevent subsequent consolidation of the memory, resulting in the observed memory deficits. On the other hand, the differential roles of the dHPC-to-RSC and dHPC-to-LS pathways involved in fear memory expression reveal either functional disassociation or compensation of multiple memory pathways. These findings convey the importance of further exploration on the neural mechanisms of these pathways to better understand functional neuroanatomy for potential clinical application.

## Methods

### Mice

Male C57BL/6J mice (n = 131; 25–32 g; 10–13 weeks old at the time of surgery) purchased from Jackson Laboratories were used for behavioral tests. After surgery, mice were dually housed (except for optrode mice that were singly housed; Figure 5) in standard mouse cages (25 × 15 × 15 cm) that contained bedding and environmental enrichment (cotton and wood sticks) on a 12 hr light/dark cycle with *ad libitum* access to water and food. Mice were removed from the study if the optical fiber headcap fell off before or during experiments (n = 3). All experimental procedures were approved by and in accordance with the Institutional Animal Care and Use Committee at Drexel University College of Medicine.

### Viral Vectors

Addgene or the University of Pennsylvania Penn Vector Core produced the adeno-associated virus serotype-1 (AAV1) encoding halorhodopsin (eNpHR3.0), enhanced yellow fluorescent protein (eYFP) or enhanced green fluorescent protein (eGFP): AAV1.CaMKIIa.eNpHR3.0.eYFP.WPRE.hGH (Addgene, #26971), AAV1.CaMKII.eYFP. WPRE.hGH (Addgene, #105622), AAV1.Syn.eGFP.WPRE.bGH (Penn Vector Core, #CS1221), respectively. The final viral concentrations were 2.66 × 10^13 GC (genome copies)/mL, 1.00 × 10^13 GC/mL and 2.80 × 10^13 GC/mL, respectively.

### Stereotaxic surgery

For optogenetic experiments, pairs of mice were randomly assigned to either experimental (Halo) or control (Ctrl) groups and received the corresponding bilateral AAV viral injections into the dHPC (0.2 µL per injection site; Halo group: AAV1.CaMKII.eNpHR3.0.eYFP; Ctrl group: AAV1.CaMKII.eYFP or AAV1.Syn.eGFP for a small group of mice). Mice were anesthetized by ketamine/xylazine solution (∼100/10 mg/kg, i.p., Vedco, Inc.) and placed into a Kopf stereotaxic instrument. Breathing was monitored throughout the duration of the surgery to ensure that anesthesia was maintained. After leveling Bregma with Lambda, three ∼340-µm holes were drilled: two bilaterally above the dHPC and one above the midline of either the RSC or LS, depending on experimental group. After drilling, viruses (200 nL) were microinjected into the dHPC by a syringe pump (World Precision Instruments, WPI) over four minutes (50 nL/min), with an addition of five minutes before removal of the injection needle (34 gauge, beveled; WPI). The coordinates for the dHPC viral injections were AP -2.0 mm, ML ±0.9 mm, DV -1.6 mm. Mice also received an optical fiber (200-µm diameter; ThorLabs, Inc.) implanted into either the RSC or LS that was secured with biocompatible ionomer (DenMat Geristore). The coordinates for the RSC optical fiber placement were AP -1.5 mm, ML 0.0 mm, DV -0.7 mm; the coordinates for the LS optical fiber placement were AP 0.6 mm, ML 0.0 mm, DV -2.05 mm. Behavioral experiments occurred four-five weeks after surgery to allow gene expression.

For optogenetics combined with electrophysiology recording, an additional four mice were used. These mice received AAV injections into the dHPC (0.2 µL; AAV1.CaMKII.eNpHR3.0.eYFP) with the same dHPC coordinates as above. Mice underwent a similar surgical procedure as above, except an optrode, an optical fiber attached to a bundle of eight tetrodes, was implanted into either the RSC (n = 2) or LS (n = 2). This optrode was coupled with a microdrive to gradually drive the electrodes to deeper recording sites post-surgery, as seen in our previous publications (Wang et al., 2015, Opalka et al., 2020). The coordinates for the RSC and LS were slightly adjusted more laterally in order to record cell bodies: the coordinates for the RSC optrode placement were AP - 1.5 mm, ML 0.3 mm, DV -0.9, while the LS optrode placement were AP 0.6 mm, ML 0.2, DV -2.0 mm. Recording occurred at least three weeks after surgery to allow gene expression of eNpHR3.0 on hippocampal terminals.

### Experimental Design

Four-five weeks after surgery, mice received two days of handling, 10–15 min on the first day and ∼5 min on the second day. After handling, mice were trained with a contextual fear conditioning procedure in four identical footshock chambers (32 × 25 × 25 cm) inside sound-attenuating cubicles (64 × 75 × 36 cm; Med Associates, Inc.), silencing the dHPC projection either during acquisition or retrieval of recent and remote contextual memory. To reduce the mouse’s time spent in the footshock chamber before recording began, two chambers were used when one experimenter was present, or four chambers were used when two experimenters were present. All behavioral procedures began around 4:00 PM. During acquisition and retrieval tests, mice were counterbalanced among pairs and tested in separate chambers. After each mouse was tested, the chamber was cleaned with 70% ethanol.

### Optogenetic Inhibition during Contextual Fear Acquisition

During acquisition (day 0), the optical fiber implant of each mouse was connected to a 532 nanometer (nm) green laser (Opto Engine LLC). Then, mice were placed into the footshock chamber (Med Associates, Inc.) and allowed to explore for a total of 270 seconds (s), receiving laser stimulation throughout the entire duration. Three footshocks (2 s, 0.75 mA) were delivered at 120 s, 180 s, and 240 s. After acquisition, mice were tested for recent memory the next day (day 1) and remote memory in a month (day 31). Each memory test included a 300 s exploration period in the same footshock chamber as day 0 but with no laser stimulation. Videos of behavior and freezing scores were collected utilizing video-tracking software (VideoFreeze; Med Associates, Inc.) to determine freezing durations amongst groups. For laser stimulation, we used a power of ∼6 milliwatts (mW) throughout experiments except for a subset of mice used in Figure 1 in which two RSC datasets of mice were tested (24 per set; n = 12 per group). The first set received ∼6 mW laser stimulation, whereas the second set received ∼0.3 mW laser stimulation (to minimize any leakage of laser into hippocampal regions since the RSC is anatomically close to the dHPC). Despite the large difference of the two laser powers used, the results from the two groups were similar (Suppl. figure 3); thus, we combined the two datasets for analysis (Figure 1).

### Optogenetic Inhibition during Contextual Fear Retrieval

During acquisition (day 0), the optical fiber implant of each mouse was connected to a 532 nm green laser. Then, mice were placed into the footshock chamber (Med Associates, Inc.) and allowed to explore for a total of 270 s, receiving no laser stimulation. Three footshocks (2 s, 0.75 mA) were delivered at 120 s, 180 s, and 240 s. After acquisition, mice were tested for recent memory on the next day (day 1) and remote memory in a month (day 31). For each memory test, the optical fiber implant of each mouse was connected to a 532 nm green laser. Mice were then placed in the same footshock chamber as day 0 for a total of 480 s, receiving laser stimulation for the first 300 s. Data from 300–480 s are shown in Suppl. figure 5. Videos of behavior and freezing scores were collected utilizing video-tracking software (VideoFreeze; Med Associates, Inc.) to determine freezing durations amongst groups. For laser stimulation, we used a power of ∼6 mW throughout.

### Data Analysis

For behavioral experiments without the optical fiber connected to the mouse (contextual fear acquisition memory tests), freezing was calculated using the VideoFreeze software (motion index ≤ 18 and lasted for at least 1 s was considered freezing) (Anagnostaras et al., 2010, Wang et al., 2015). Freezing scores were exported from Video Freeze, and then percentage of freezing per minute was further calculated in MATLAB. For behavioral experiments with the optical fiber connected to the mouse (all training and contextual fear retrieval memory tests), we noticed that sometimes mice were visibly freezing, but the VideoFreeze software calculated swaying of the optical fiber tether as mouse movement, rendering the freezing calculation inaccurate. Thus, we reduced this fiber movement effect by cropping the top 75% of video frames that almost exclusively recorded fiber but not mouse movement (mice were typically recorded in the bottom 25% of video frames by a side-view camera except during footshocks). We then used MATLAB to analyze the cut video (bottom 25%) and calculate mouse freezing index with the same parameters as VideoFreeze (motion index ≤ 18 and lasted for at least 1 s). Repeated measures two-way ANOVAs and Bonferroni post hoc tests were used for analyses. All statistics were run in SPSS (IBM). All mice were included in the analysis (n = 128) except those with their headcap detached before or during experiments (n = 3).

### In vivo electrophysiology

We used optrodes, optic fibers attached to eight tetrodes for recording, similar to that shown previously (Wang et al., 2015, Opalka et al., 2020). Each tetrode consisted of eight wires (90% platinum and 10% iridium; 18 μm diameter; California Fine Wire). About two weeks after surgery, tetrodes were screened for neural activity. Neural signals were preamplified, digitized, and recorded using a Neuralynx Digital Lynx acquisition system. Each electrode bundle was lowered by 50−100 μm daily until multiple RSC or LS neurons were detected. Spikes were digitized at 30 kHz and filtered at 600–6000 Hz, using ground as the reference for both. All mice received 2–5 sessions of recording when they were freely-behaving or sleeping in home cages (∼2 h). After completion of each recording session, the electrode bundle was lowered by 50−100 μm to record a deeper site in the RSC or LS: a total of 2–5 sites were recorded from each mouse. 5-min continuous laser stimulations (∼6 mW; 532 nm; Opto Engine LLC) were used in each session (∼2 h), with an inter-trial interval of 15 min.

### Histology

After completion of remote memory tests (or locomotor tests when applicable), mice received pentobarbital overdose and were perfused transcardially with 1× PBS for 4 minutes followed by 10% formalin at a rate of 1.3 mL/min until muscle contractions occurred. Then, 10% formalin was perfused for 30 min at a rate of 0.3 mL/min. Brains were stored in 10% formalin for at least 24 hours before vibratome sectioning (Leica VT1000 S). Coronal sections (50 µm) were collected to verify all injection sites, viral expression, and optical fiber placements.

## Acknowledgements

We thank the Histology and Imaging Facilities at Drexel University Department of Neurobiology and Anatomy for their generous help. We also thank Drs. Jed Shumsky and Rodrigo España for advice on statistics and Candace Rizzi-Wise for the editing of our manuscript. This research was supported by Drexel University College of Medicine and NIMH/NIH (R01 MH119102).

## Author Contributions

ANO and DVW designed and conducted the experiments, analyzed the data, and wrote the manuscript.

## Competing interests

The authors declare no competing interests.

## Data availability

The datasets generated and/or analyzed during the current study are available from the corresponding author on request.

## Supplementary Methods

### Locomotor Test

After remote memory tests, a subset of mice from the LS Acquisition (n = 7; 3 Ctrl and 4 Halo) and LS Retrieval (n = 8; 4 Ctrl and 4 Halo) experiments received a locomotor test (day 36) for 600 s, receiving laser stimulation for the first 300 s. The optical fiber implant of each mouse was connected to a 532 nm green laser. Then, mice were placed into the same footshock chamber, instead, with a cyclic wall, smooth floor, lights off in both the testing chamber and room, and 1% acetic acid scent. (This modification created a new context, which induced minimal or no freezing). Videos of behavior and motion index were collected utilizing video-tracking software (VideoFreeze; Med Associates, Inc.) to determine total locomotor activity amongst groups. Motion index was normalized by a ratio of individual motion index to average control motion index. For laser stimulation, we used a power of ∼6 mW throughout.

**Suppl. figure 1.**
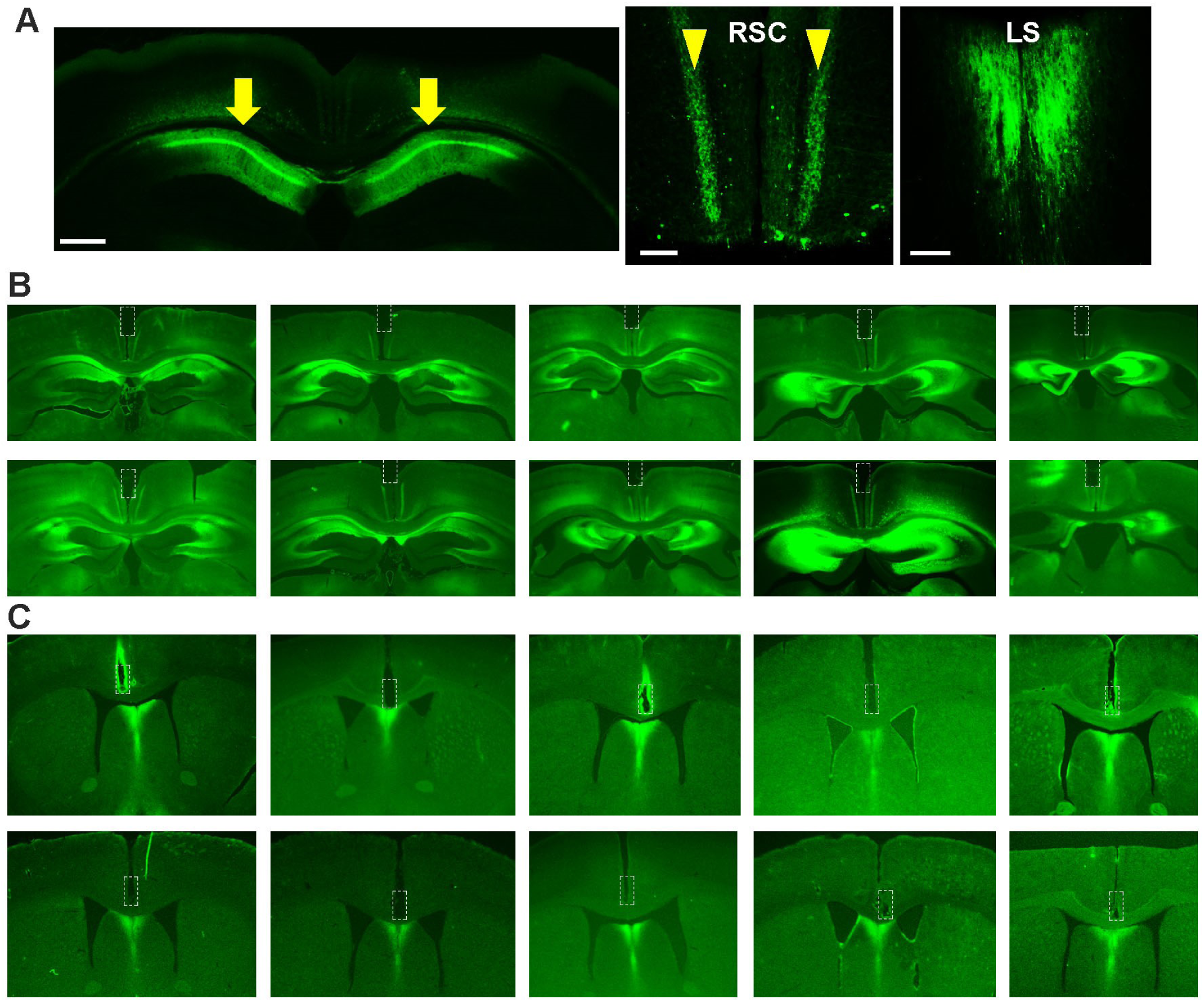
dHPC projects directly to the RSC and LS. (A) Representative coronal sections showing the injection of AAV1.CaMKII.eYFP primarily in the medial dorsal dHPC (left) and projections in granular RSC layer 3 (middle; arrows) and midline LS (right). Scale bars, 0.5 mm (left); 0.1 mm (middle & right). (B&C) Examples of optic fiber tracks (dashed squares) in the RSC (B; n = 10) and LS (C; n = 10) of mice that were bilaterally injected with AAV1.CaMKII.eNpHR3.0.eYFP into the medial dorsal dHPC.

**Suppl. figure 2.**
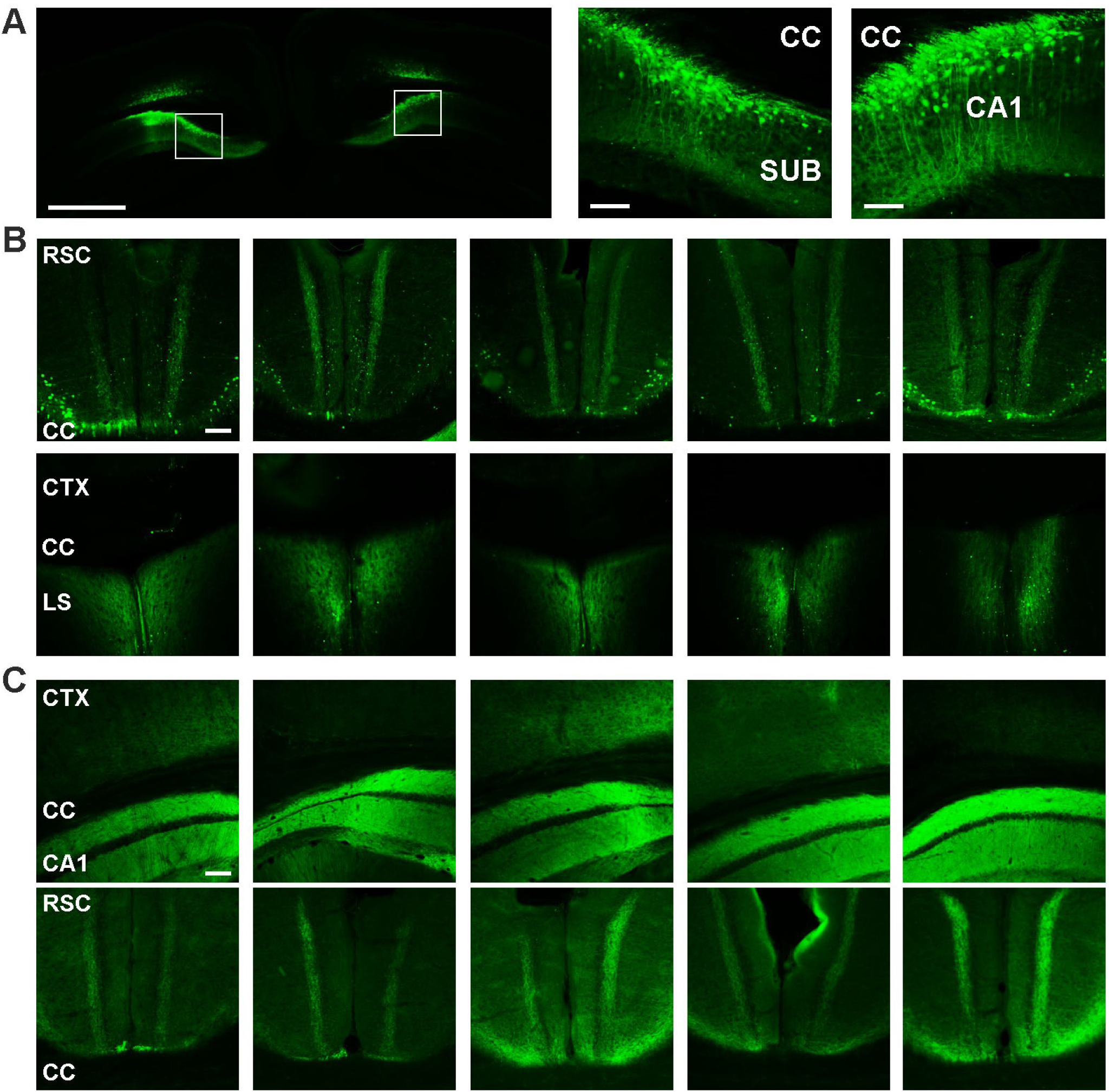
AAV viral spread. (A) Representative coronal sections of dHPC showing the spread of AAV1.CaMKII.eYFP to dorsal subiculum (SUB; left inset) and caudal dorsal CA1 (right inset). (B) Examples of partial spread of AAV1.CaMKII.eYFP (n = 5 mice) in the ventral lateral RSC near corpus callosum (CC; top panels), but not in the LS (bottom panels). (C) Examples of partial spread of AAV1.CaMKII.eNpHR3.0.eYFP (n = 5 mice) to cortex (CTX) dorsal to CA1 and RSC (bottom panels). Scale bars: 0.1 mm except A (top left), which is 1.0 mm.

**Suppl. figure 3:**
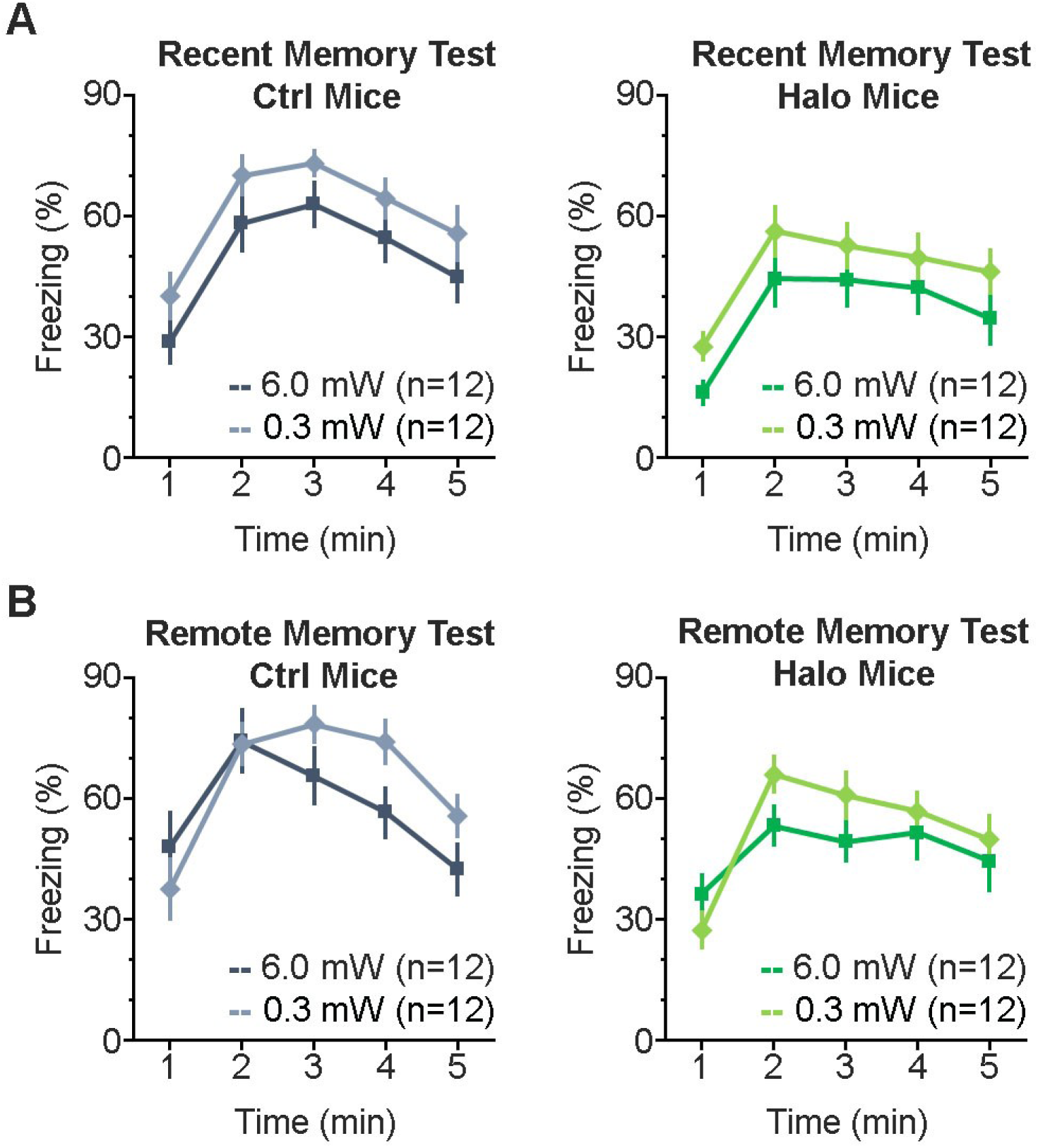
Different power of laser stimulation of the dHPC-to-RSC pathway during training had similar effect on freezing during memory tests. (A) Percentage of freezing across five minutes during the recent memory test (day 1). There is no significant difference of freezing (mean of 5 minutes) between laser power for Ctrl mice (left, *t*_(22)_ = -1.637, *P* = 0.116) and Halo mice (right, *t*_(22)_ = -1.425, *P* = 0.168). (B) Percentage of freezing across five minutes during remote memory tests (day 31). There is no significant difference of freezing (mean of 5 minutes) between laser power for Ctrl mice (left, *t*_(22)_ = -0.893, *P* = 0.382) and Halo mice (right, *t*_(22)_ = -0.833, *P* = 0.414). Error bars indicate SEM.

**Suppl. figure 4:**
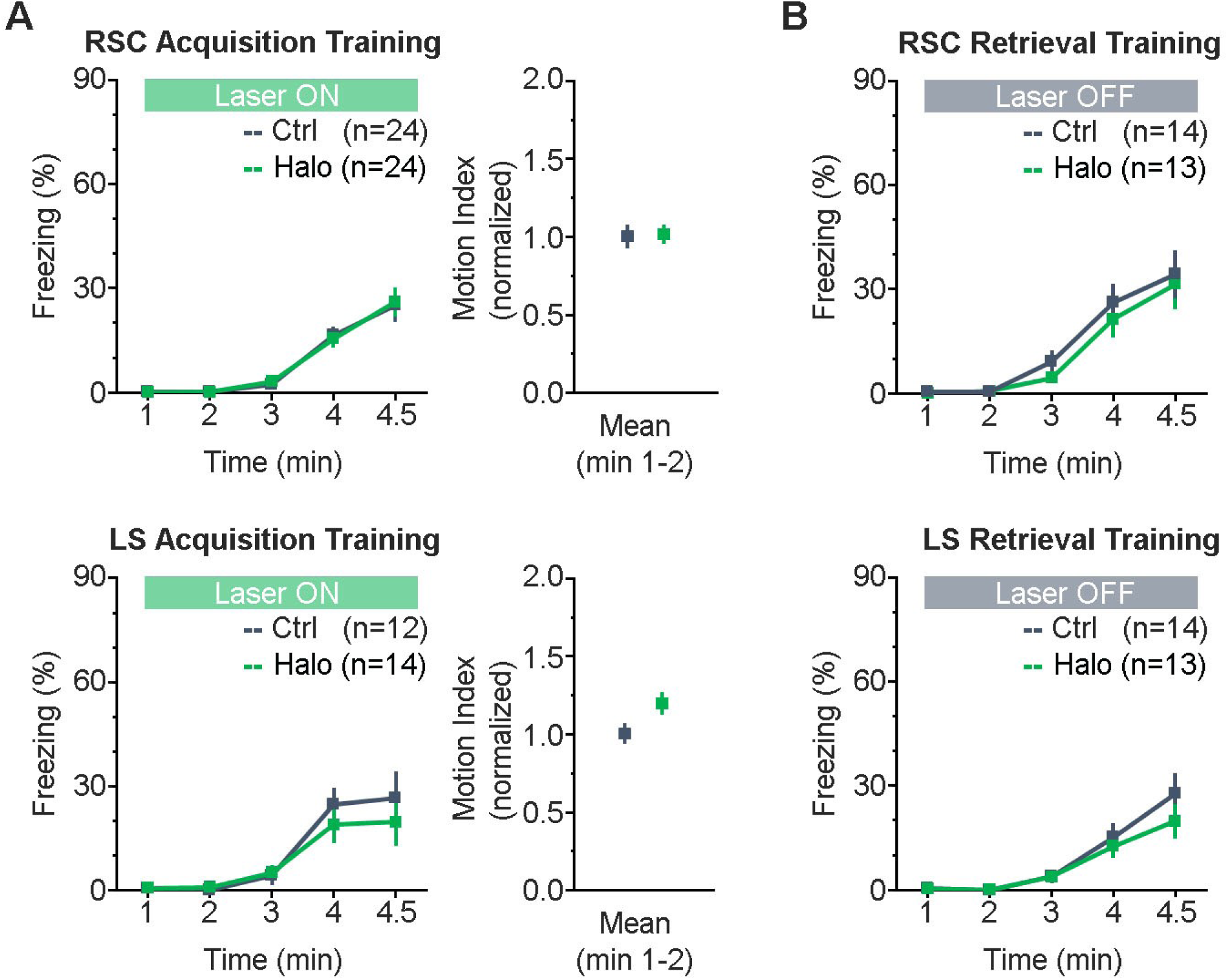
Laser stimulation has no effect on freezing during training. (A) Left panels, percentage of freezing across four and a half minutes during acquisition (day 0) revealed no significant difference between Ctrl or Halo groups during laser stimulation of dHPC projections to the RSC (top) or LS (bottom). Right panels, normalized motion index during the first two minutes (before the onset of footshock) for the RSC (top) and LS (bottom) groups. (B) Percentage of freezing across four and a half minutes during acquisition (day 0) revealed no significant difference between Ctrl or Halo groups without laser stimulation of dHPC projections to the RSC (top) or LS (bottom). Error bars indicate SEM. *P* > 0.05 for all comparisons (two-tailed unpaired *t* test).

**Suppl. figure 5.**
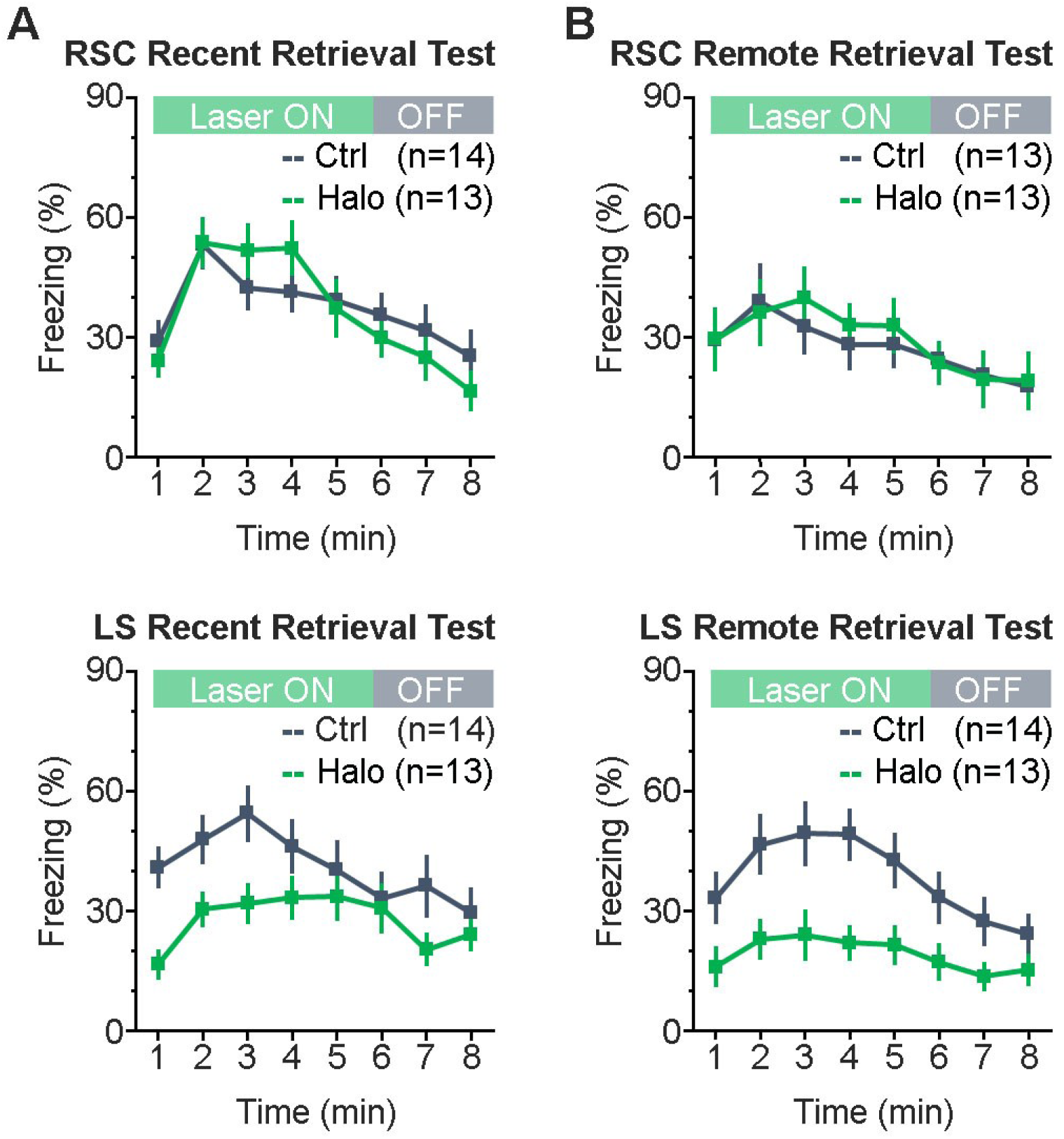
Memory retrieval with and without laser stimulation. Minute-by-minute percentage of freezing for the dHPC-to-RSC (top) and dHPC-to-LS (bottom) mice during the recent (A) and remote (B) contextual fear memory tests, respectively. Note that there’s no increase of freezing during minutes 6–8 (laser off) compared to minutes 1–5 (laser on).

